# The geometry of prey capture in praying mantis forelegs

**DOI:** 10.1101/2024.04.18.590067

**Authors:** Shu D. Dan, Danielle S. Taylor, Jaime Yockey, Gavin J. Svenson, Joshua P. Martin

## Abstract

The form of an animal’s limbs has to balance multiple functions: locomotion, grasping, climbing, and jumping, among others. For cryptic animals, especially those that resemble elements of their habitat like sticks or grasses, the limbs may also be modified to enhance the camouflage. The performance of a limb in one category may require a tradeoff, reducing performance in another category. Praying mantises provide a diverse group of insects who all use their forelegs for one function, capturing prey, while some species use them as part of their camouflage. Here we use a large database of images of mantis species to capture the variation in morphology across the order, and to calculate the largest prey that their forelegs can hold. We find that the length and thickness of the femur and the length of the tibia comprise most of the variability across species. The majority of species have similar foreleg morphology, with two large groups extending into areas of the morphospace with thicker or thinner forelegs. A geometric relationship between dimensions of the foreleg and the optimal prey diameter maps directly onto the variability across species determined by principal components analysis; legs with thinner femurs and shorter tibia can’t hold large prey, and the distribution of the species across the morphospace follows the gradient of optimum prey size. These results suggest that some species trade ability to grasp larger prey for benefits including crypsis, and the praying mantises are an ideal system for studying morphological and functional variation in limbs.

## Introduction

Evolution shapes the limbs of animals to serve multiple purposes. An insect limb may be adapted for motor functions (e.g. walking, climbing, jumping, and grasping prey), and also modified superficially for camouflage (e.g. grass resemblance). Adaptations for a particular function may require trade-offs (Agrawal et al. 2010; Wainwright 2007) that reduce the performance of another function, entailing a cost.

Praying mantises offer a unique opportunity to study trade offs in a phylogenetic context. All mantises are predators that use their specialized raptorial forelegs to strike and hold prey. Although morphologically complex, the general structure and function of the foreleg is preserved across all praying mantis species. Large muscles in the femur flex the tibia, holding prey against the spiny surface of the femur (Levereault 1938, Prete 1990).

Adaptations for crypsis have produced an incredible diversity of body and foreleg forms in this order, yet all species depend on the foreleg for both prey capture and locomotion. In some species, raptorial forelegs exhibit extreme cryptic characteristics associated with ecomorphic specialization (Svenson and Whiting 2009) that may reduce the ability of the leg to grasp prey. The raptorial forelegs in many species are morphologically specialized to resemble elements of their environments, e.g. grasslands, tree canopies, tree trunks, and deserts (Edmunds and Brunner 1999), as a defense against predators (Edmunds 1976). For example, the grassland species *Pyrgomantis jonesi* features a conical cranial projection and abducted forelegs that fit seamlessly against the body, thereby resembling a blade of grass (Edmunds 1976, Edmunds 1990, Robinson 1973). This combination of habitat and morphological specialization is defined as an ecomorph. The legs of mantises come in a remarkable diversity of forms, and a prime example of a many-to-one mapping of morphological form to biomechanical function: while a diversity of phenotypes have arisen (e.g. ecologically disparate foreleg crypsis), the primary function (prey capture) is preserved (Alfaro et al. 2005, McGee and Wainwright 2012, Wainwright et al. 2005). However, these modifications for crypsis might compromise prey capture.

To compare the ability to capture prey across species, we need a measure that can be applied to all species equally. A simple, geometric relationship between measurements of the mantis foreleg defines the largest prey that the leg can hold, and is correlated with the preferred size of prey that it will strike (Holling 1964). Here, we use a database of digitized images of praying mantises, a machine-learning method to identify anatomical markers across diverse species, and morphological analysis to explore how the capacity to hold prey varies across mantis species.

## Results

We used a large database of digitized images of the mantis collection at the Cleveland Museum of Natural History. These mantises are all pinned using a standardized method, with their forelegs spread laterally in a plane parallel to the body, bent at approximately ninety degree angles at the coxa-femur and femur-tibia joints (Brannoch et al. 2017). This creates a consistent, two-dimensional image of the forelegs across all species necessary to accurately capture the dimensions in a comparable way for all specimens (Fig. 1C). This sample contains species in 11 of the 16 superfamilies in this order (Schwartz and Roy, 2019), and examples of major ecomorphs (Fig. 1A,B). The orientation of the leg in these images parallels the geometric measurements of ideal prey size (Fig. 1D, after Holling 1964), i.e. the largest prey, assuming an ideal spherical size, that can be held between three points on the tibia, femur, and the tip of the tibial hook. Anything larger could escape through the gap between the tibial hook and the femur, and anything smaller would provide less food for the mantis’s effort.

**Figure 1.**
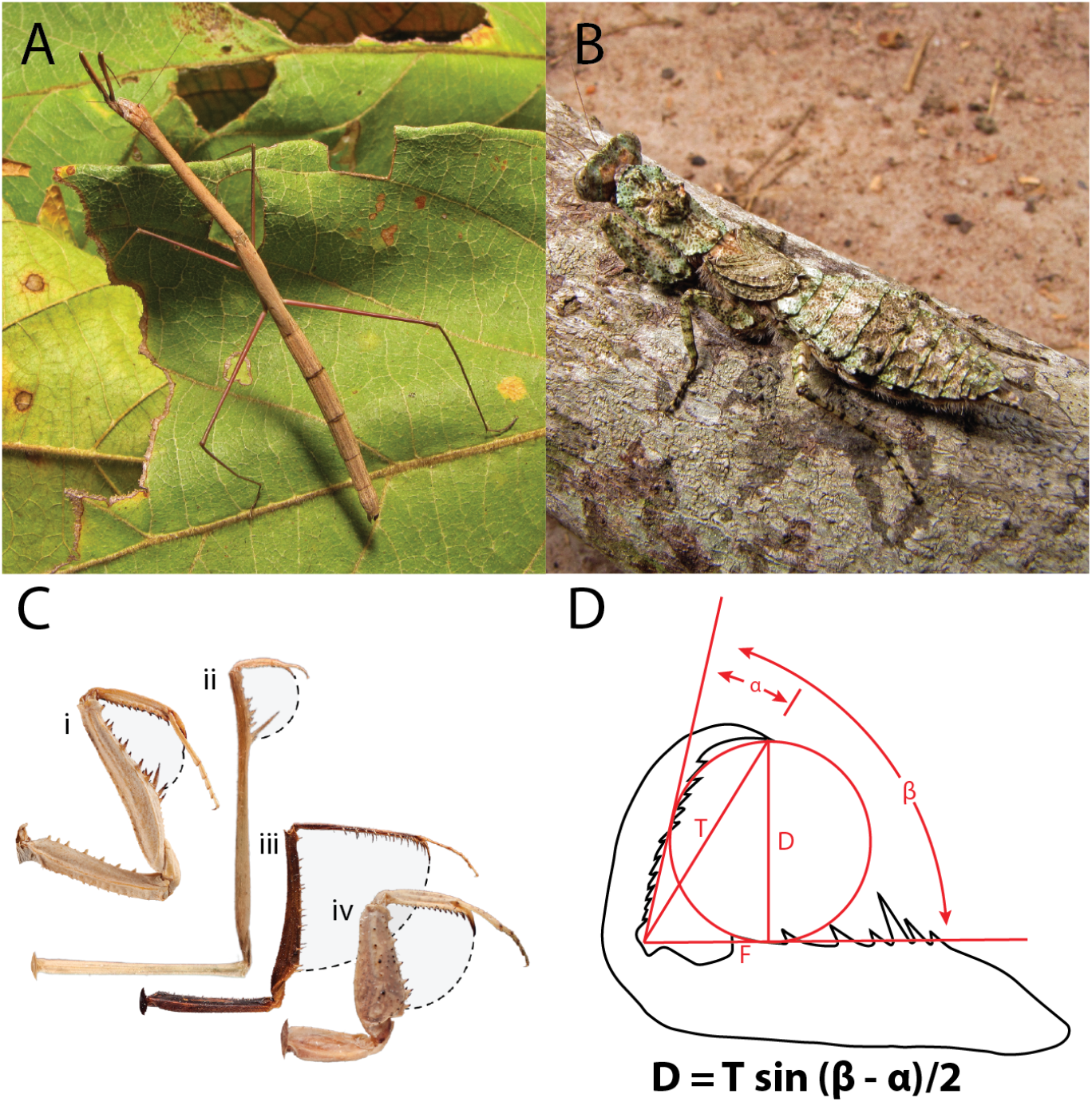
Mantis forelegs vary across species and ecomorphs. **A)** *Compsothespis sp*., an example of a mantis with adaptations for crypsis among sticks and grasses. **B)** *Tarachodes santus*, an example of a species with adaptations for crypsis on bark. **C)** Examples of the variety of foreleg forms in one group of mantises, the Tarachodinae; i - *Eremoplana iufelix*, ii - *Schizocephala bicornis*, iii - Toxoderopsis taurus, and iv - Tarachodes afzelli. **D)** The optimum (largest) prey diameter **(D)** can be calculated from the length of the tibia from the femoral-tibial joint (T), the angle of the tibial hook (a), and the optimum opening angle (β). (after Holling 1964).

We used a machine-learning approach designed for tracking anatomical features between frames of a video (DeepLabCut, Mathis et al. 2018), and adapted it to identify the same anatomical features across species of mantises. We identified 16 anatomical markers that could be consistently identified in all species (Fig. 2A). The landmarks were transformed into a common dimension using Procrustes superimposition (Fig. 2B), to scale and align all species in the same coordinate plane. Principal components were generated to reduce the 16 dimensions into two that captured 48% and 16% of the variability between species. We then used a back-transform method (Olsen 2017) to produce a hypothetical shape at a grid of points along the first and second principal component axes (Fig. 2D). Principal component 1 had the greatest correlated variability among these back-transformed points at the proximal portion of the femur, and the tibial hook (Fig 2Ci). This represents both the relative length and thickness of the femur, and the length of the tibia. Principal component 2 captured variability primarily in the thickness of the proximal femur, as well.

**Figure 2.**
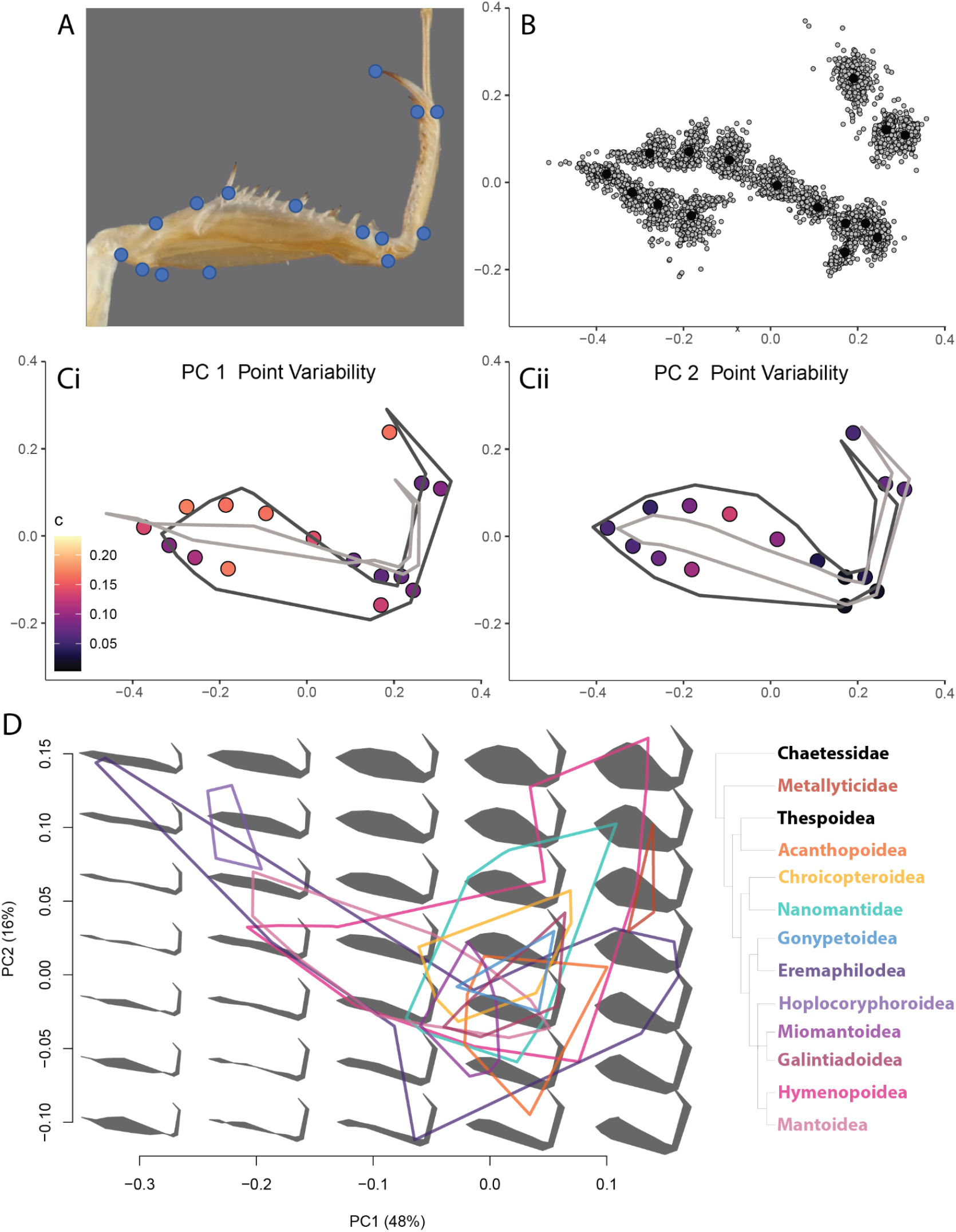
Morphometric analysis of forelegs. **A)** Example image of a mantis foreleg with the 15 points used in the morphometric analyses. **B)** Projection of points from all species included in this analysis in Procrustes space. **C)** The range of variability in Procrustes space along the first (i) and second (ii) principal component axes. Points are the median back-transformed morphology, color-coded by the amount of variability (Euclidean distance) at each point along this PC axis. Lines are the back-transformed morphology of the shapes at the minimum (light gray) and maximum (dark gray) ends of each PC axis. **D)** Morphospace along the first two principal components. Gray shapes are the back-transformed shape at evenly spaced points in this space. Lines are a contour surrounding the individual species in the Superfamilies listed on the left.

We mapped the phylogeny of mantises to this “morphospace” defined by the first and second principal components (Fig. 2D). The majority of superfamilies, including all but one (Metallyticidae) of the most basal groups, occupy a median location in the morphospace. The Metallyticidae were an outlier in this regard, separated from the other groups on both axes, with generally shorter and fatter forelegs. Several superfamilies (Acanthopoidea, Chroicopteroidea, Gonypetoidea, Miomantoidea, and Galintiadoidea) overlapped in a narrow range of variability on both axes. The others were larger groups that were represented both in that central area, as well as two offshoots into the upper right and upper left quadrants of the morphospace. Some Nanomantidae species were represented in that upper right quadrant of thicker, squatter forelegs, while members of the Mantoidea extended into the upper left quadrant of longer, thinner forelegs with relatively shorter tibia. The large superfamilies of Hymenopoidea and Eremaphilodea occupied all three regions. Finally, species in the Hoplocoryphoroidea were found only in the upper left quadrant, with only longer and thinner forelegs. We also note that there are regions of the morphospace, the lower left quadrant in which no species are found; that is, there are no species whose legs have those shapes.

We turn next to the key measurements necessary to calculate the optimal prey diameter (from Fig. 1D); tibial length, tibial hook angle, and optimal opening angle, and the result of that calculation: optimum prey diameter. These were calculated from the Procrustes projection of the anatomical markers on the legs, to normalize the scale across species. The first two, tibial length and tibial hook angle, are measured directly from the markers. Optimal opening angle is calculated from a geometrical relationship between those values (Holling 1964, Holling et al. 1979). The largest variability between groups captured here is the shorter tibial lengths of the Hoplocoryphoroidea (Figure 3A). That group overlaps more with the others in tibial hook angle and opening angle (Fig. 3B,C), but has the largest number of species with smaller optimal prey diameter (Fig. 3,D). Groups that occupied the upper left quadrant followed a similar pattern among the minority of species in that quadrant. There was no clear pattern among these measurements for the legs that occupied the upper right quadrant. The peaks of the optimum prey diameter density distributions cluster around the population mean, with only a few groups (Eremaphiloidea, Gonypetoidea) skewed towards larger prey diameter (Fig. 3D).

**Figure 3.**
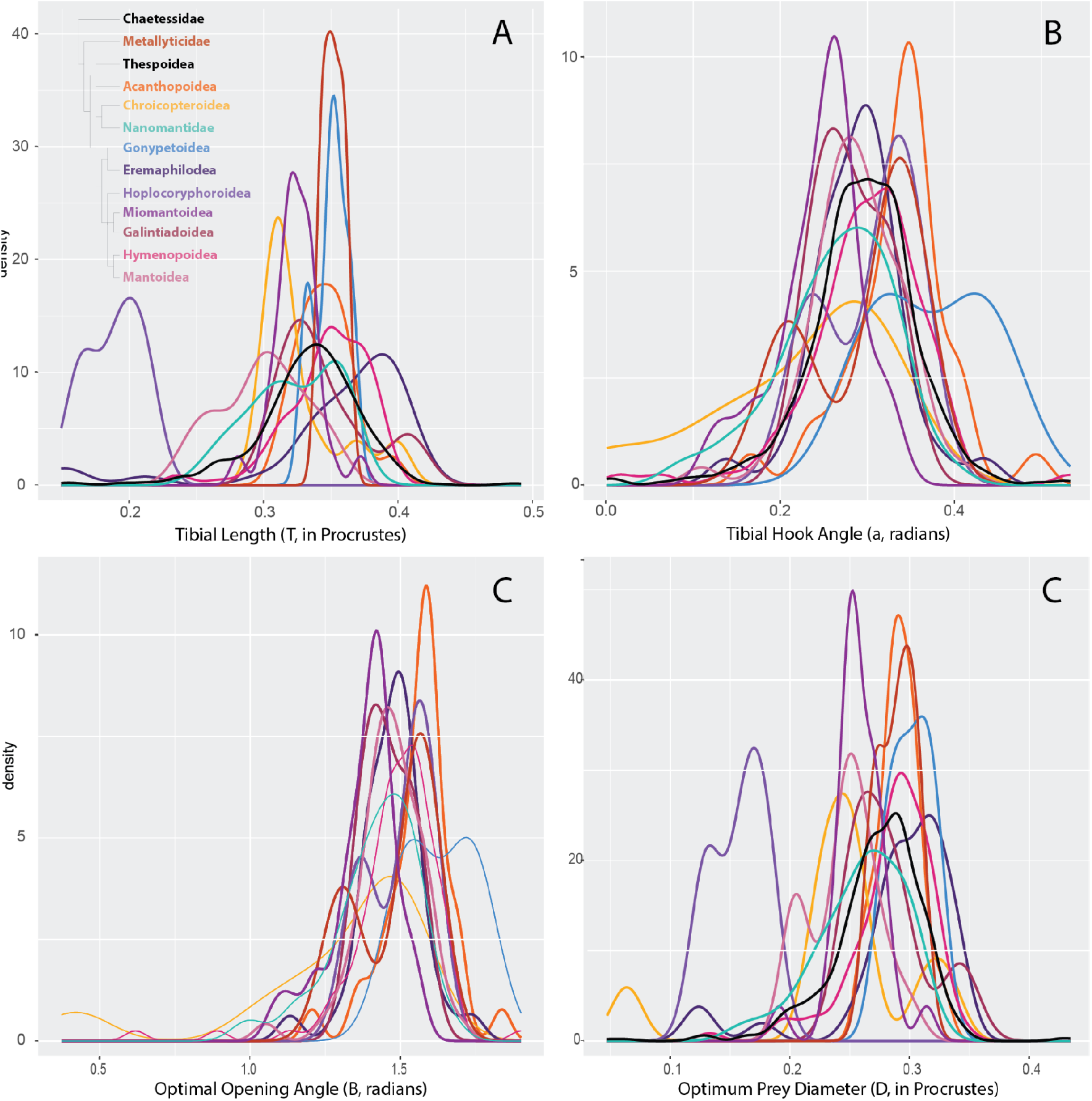
Variability in foreleg dimensions among mantis clades. Histograms of the **A)** tibial lengths, **B)** tibial hook angles, **C)** optimal opening angle, and D) optimum prey diameter for the mantis Superfamilies (inset, upper left) and the whole population of mantises (black line).

The back-transform method allows us to calculate geometrical features of not only the real forelegs, but also the hypothetical, back-transformed shapes that occupy all areas of the morphospace. We first note a trend in the optimum prey diameter calculated for each species (Figure 4A). Forelegs toward the right along the PC1 axis can hold increasingly larger prey.

**Figure 4.**
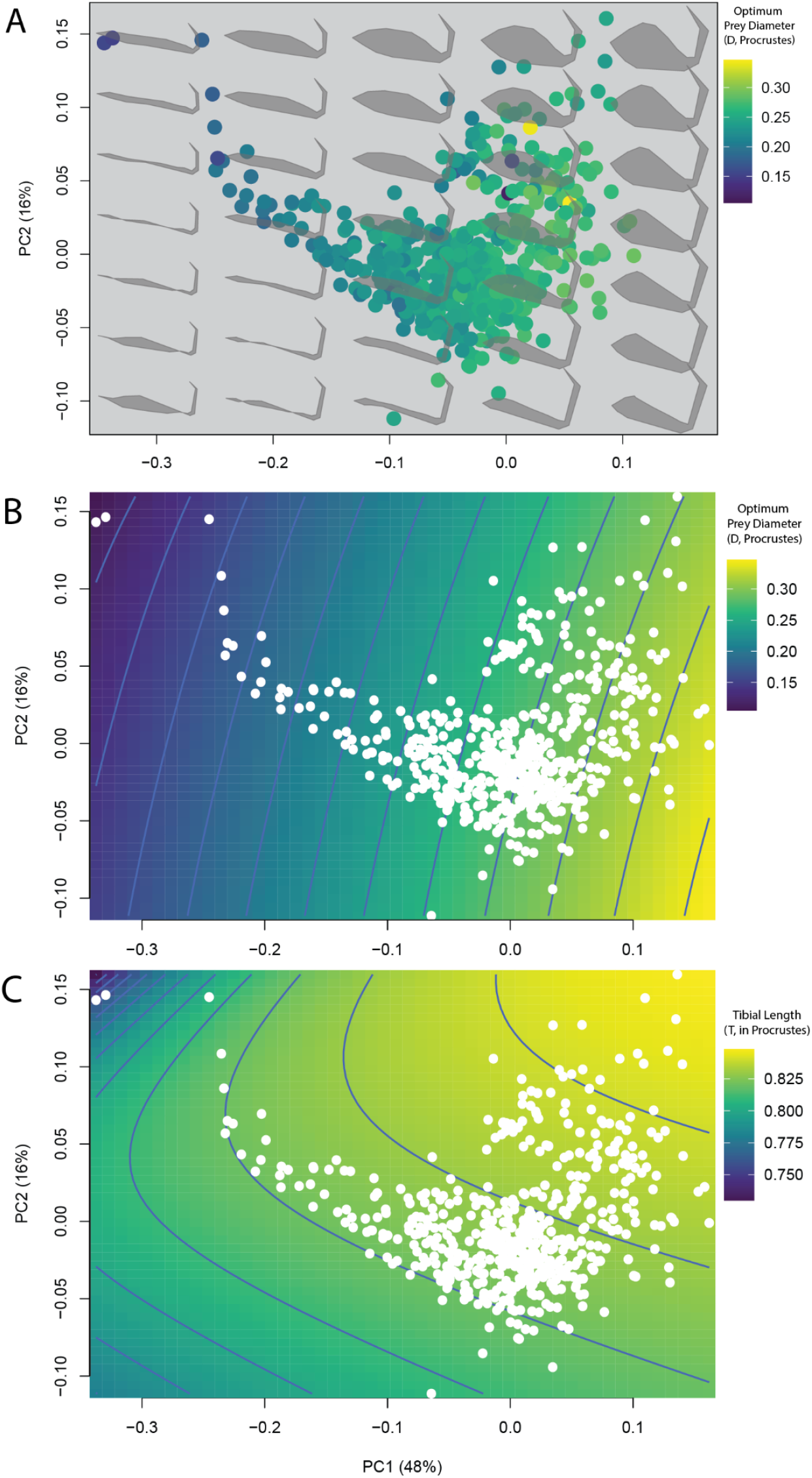
Foreleg dimensions across PC morphospace. **A)** Morphospace from figure 2D, with optimum prey diameter calculated for each species (color scale). **B)** Morphospace with calculated values of optimum prey diameter for back-transformed foreleg morphologies. **C)** Morphospace with calculated values for tibial length for back-transformed foreleg morphologies.

Next we use the back-transformed marker points to measure and calculate the variables of the geometric model. We divide the morphospace into a dense grid, produce a back-transformed shape at each point, and calculate the optimum prey diameter. Remarkably, the gradient of the optimum prey diameter aligns well with the spread of the real species across the morphospace (Fig. 4B). Species towards the lower right quadrant, where the majority of superfamilies overlap (Fig. 2D), have a larger optimum prey diameter. The tail of the cluster of species leading to the upper left quadrant follows a gradient of lower prey diameter, suggesting that species that have evolved longer, thinner legs have lost some ability to grasp larger prey.

Tibial length also maps to the distribution of species in the morphospace (Fig. 4C). The legs in the upper right climb a gradient towards a relatively larger tibia. The majority of species in the central, median cluster follow along a gradient line towards the upper left quadrant, and the furthest outlying species in the upper left quadrant fall down the gradient towards shorter tibia. Although the morphospace is defined by a process that captures variability across all anatomical markers, it is clear from these results that the geometry of prey capture maps directly onto the morphospace, and therefore is likely a key driver of that variability.

## Conclusions

While the forelegs of praying mantises come in a large variety of forms, they are all used to capture prey in a characteristic rapid strike. The present results demonstrate that some mantises have forelegs with reduced ability to capture prey. Critically, the morphospace was calculated naively and independently of the geometric analysis, yet the optimum prey diameter maps directly onto the largest axis of variability. This suggests that the species in this tail of long, thin forelegs have sacrificed some ability to capture prey, likely for the benefit of cryptic forms that require thin appendages.

The majority of species and taxonomic groups have a similar ability to capture larger prey, while four of the superfamilies examined had one or more species with reduced capacity to capture prey. Of these, three had the majority of species with capacities near the median, and in only one superfamily (Hoplocoryphoidae) all of the examined species had reduced capacity. This was driven primarily by having a relatively short tibia; other factors in the geometric analysis were more similar between species. While the habitat of many of these species has not been reported, those that are known are found in grasses or brush, and have apparently evolved special resemblance to that habitat that compromised their ability to capture prey.

While biomechanical adaptations can increase diversity by allowing species to exploit new resources (Alfaro et al. 2005; McGee and Wainwright 2012; Wainwright et al. 2005, Price et al. 2012; Herrel et al. 2005), recent work is beginning to examine how trade-offs may limit diversity across groups of similarly adapted organisms (Holzman et al. 2012; Ghalambor et al. 2004). Our data set was determined by the species that had been digitized, and among those only the specimens pinned so that the foreleg was parallel to the lens of the camera. An analysis of diversity would require a more complete, representative sample of species across groups. However, our data suggest that the reduced ability to capture prey may limit the diversity of species in those lineages, potentially requiring a trade-off between prey capture and crypsis.

The morphological analysis also suggests other biomechanical limitations among some species. The first principal component captures variability in the width of the femur. This is one of the dimensions that determines the volume of the flexor muscle, which provides the force to close against struggling prey. The species that occupy the upper right quadrant of the morphospace have larger and thicker femurs, yet the geometric relationship shows that they cannot grasp larger prey than the majority of the other species. Adaptation in this dimension may allow for a larger volume of flexor muscles, to hold struggling prey. Many of these species are of the bark and ground-mantis ecomorphs, which run on flat surfaces and do not generally resemble sticks or grasses.

Biomechanical analysis can use anatomical features of muscle and skeleton to predict the performance of limbs (Goyens et al. 2014; van der Meijden et al. 2012; Heethof and Norton 2009; Claverie et al. 2010; Patek et al. 2004; Blanke et al. 2017], and this initial report suggests that a model that takes into account the muscle volume could reveal further deficits, gains, or trade-offs among some species. Additionally, the forelegs are also used for locomotion - mantises that live in dense, three-dimensional environments and climb through them to find prey would have different requirements than their relatives that primarily run on flat surfaces. Mantises can be classified by their primary hunting strategy: either ambush, pursuit, or generalist species that use both (Svenson and Whiting 2009). This is another factor that is likely shaping the evolution of the foreleg in different species. Many of the pursuit species are found in the upper right quadrant of the morphospace, while many of the ambush specialists are found in the upper left.

This analysis is limited to a single geometric relationship between features of the foreleg and the largest prey that the foreleg can hold. This is necessarily a simplification that allows us to compare widely across species using anatomical markers that are consistent among all of them. Other factors contribute to the ability of an insect leg to hold prey. For example, the length of the large discoidal spine near the proximal end of the femur is also highly variable (personal observation). These spines have a flexible base, creating a one-way trap for prey (Loxton and Nicholls, 1979). In addition, the present analysis assumes a perfectly spherical prey, while real prey has varied shapes and stiffness. Indeed, mantises have been recorded capturing small birds (Nyffeler et al. 2017), well beyond the size of prey predicted by their foreleg geometry. The present, simplified analysis allows us to include all mantis species in a single functional description - species-specific adaptations can increase the ability of some species, but would require a species-specific geometrical model to calculate.

Praying mantises (Insecta: Mantodea) provide an ideal model system for exploring the consequences of functional morphology and biomechanical trade-offs on evolution. These results are exemplary of many-to-one mapping of morphological form to biomechanical function (Alfaro et al. 2005; McGee and Wainwright 2012; Wainwright et al. 2005): while a diversity of phenotypes have arisen (e.g. ecologically disparate foreleg crypsis), the primary function (prey capture) is preserved.

## Methods

### Morphometric analysis

DeepLabCut was trained to label 11 major morphological landmarks on the femur identified from previous unpublished study as well as five major landmarks on the tibia (Figure 3). Five hundred and sixty-five photos of the full body of diverse species of praying mantis in dorsal view were obtained from The Mantodea Image Database at the Cleveland Museum of Natural History (Stackshot z-stepper, a Canon 5D II SLR, macro lenses (50mm, 100mm, and MP-E 65mm), and Speedlight 580EX II flash units). The photos were resized proportionally to crop out only the right forelegs to reduce image size and future processing time. The cropped image was made into a video as one photo per frame and two frames per second, which was used as the training video for DeepLabCut. The program was set to randomly select 50 frames from the video as the training set and 16 major landmarks in each frame were manually labeled on a local machine. The landmarks were chosen to capture the geometry of the mantis forelimb using external identifiable features that could be found on all species.

With the labeled frames, the neural network was trained for 50000 iterations on the Colby server. Each photo in the video was treated as five frames and resulted in 2800 frames with predicted labels. The labeled output was manually validated and frames with obviously wrong labels were refined. The landmarks were output in the form of relative x, y coordinates in the photos and a mean output for each photo was calculated.

The landmarks were scaled and aligned using Procrustes superimposition with the geomorph R package (Baken et al., 2021 & Adams et al., 2022). Then principal component analysis (PCA) was used to reduce the 16 landmarks into two principal components (PC). Additional analysis on this data was performed using the PC space (“morphospace”). For each point in the morphospace, a “back transformed” method was used to simulate the morphology of a limb that would occupy that point (Olsen 2017). We use the actual dimensions of the limbs measured from the photographs and the projected limbs in the morphospace to calculate the geometry of the limb and the optimum prey size.

### Optimum prey size calculation

The tibial-hook angle (α) is measured from the anatomical markers, and then the optimal angle between the tibia and femur (β) can be calculated by solving the equation 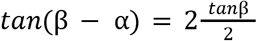 (Holling 1964, Holling et al. 1979). Then the equation *D* = *Tsin*(β - α) can be used to calculate the optimum diameter (D) of the prey, where T=the length of the tibia from the femur-tibia joint to the tip of the tibial hook. All calculations were performed using custom scripts in R (R Core Team 2021).

## Acknowledgements

This project was supported under the NSF grant DEB-1216309 to Gavin J. Svenson and the NSF grant IIS-1704366 to Joshua P. Martin.

## Author Contributions

S.D.D. and D.S.T. collected data, performed analysis, and participated in writing the manuscript.

J.Y. assisted in collecting data and performing analysis. J.P.M performed analysis, and G.J.S and J.P.M. conceived the idea, designed the study, and wrote the manuscript.

## Notes

### Competing Interest Statement

The authors have declared no competing interest.

